# Chemoresistome Mapping in Individual Breast Cancer Patients Unravels Diversity in Dynamic Transcriptional Adaptation

**DOI:** 10.1101/2023.02.09.527790

**Authors:** Maya Dadiani, Gilgi Friedlander, Gili Perry, Nora Balint-Lahat, Shlomit Gilad, Dana Morzaev-Sulzbach, Anjana Shenoy, Noa Bossel Ben-Moshe, Anya Pavlovsky, Eytan Domany, Iris Barshack, Tamar Geiger, Bella Kaufman, Einav Nili Gal-Yam

## Abstract

Emerging evidence reinforce the role of non-genetic adaptive resistance to chemotherapy, that involves rewiring of transcriptional programs in surviving tumors. We combined longitudinal transcriptomics with temporal pattern analysis to dissect patient-specific emergence of resistance in breast cancer. Matched triplets of tumor biopsies (pre-treatment, post-treatment and adjacent normal) were collected from breast cancer patients who received neo-adjuvant chemotherapy. Full transcriptome was analyzed by longitudinal pattern classification to follow patient-specific expression modulations. We found that dynamics of gene expression dictates resistance-related modulations. The results unraveled important principles in emergence of adaptive resistance: 1. Genes with resistance patterns are already dysregulated in the primary tumor, supporting a primed drug-tolerant state. 2. In each patient, multiple resistance-related genes are rewired but converge into few dysregulated modules. 3. Rewiring of diverse genes and pathway dysregulation vary among individuals who receive the same treatments. Patient-specific chemoresistome maps disclosed tumors’ acquired resistance and exposed their vulnerabilities.

Mapping the complexity of dysregulated pathways in individual patients revealed important insights on adaptive resistance mechanisms. To survive the toxic drug effect, tumor cells either sustain a drug-tolerant state or intensify it, specifically bypassing the drug’s interference. Depicting an individual road map to resistance can offer personalized therapeutic strategies.

## Introduction

Despite the considerable importance of tumor drug resistance to cancer morbidity and mortality, our comprehension of the various molecular mechanisms involved in resistance is limited [1]. The actual response of an individual patient remains a ‘black box’. The oncologist cannot accurately predict which tumor, eligible for a chemotherapy, will be eliminated by the drugs, and the intrinsic resistance to chemotherapy remains a substantial enigma. Recently, the prevailed genetic clonal selection, as the main mechanism of resistance is challenged by emerging indications of non-genetic adaptive mechanisms of drug resistance [2,3]. Cumulative evidence show that in response to treatment, tumors adopt a drug-tolerant state through transcriptional reprogramming [4–6]. To survive chemotherapy, cancer cells rewire molecular pathways, thereby, escaping anti-proliferative drugs. However, as cancer is a continuous process, the individual route to resistance in each cancer type and each patient is elusive.

Breast cancer is currently the most common type of female cancer and the second cause of cancer mortality in women[7]. Major international efforts have profiled primary tumors and defined the molecular architecture of breast cancer [8–12]. It is now clear that breast cancer is a heterogeneous disease, classified into distinct subtypes: hormone receptor positive (HR+), HER2 positive and triple negative breast cancer (TNBC). While novel targeted therapies and immunotherapies are being developed, chemotherapy remains a mainstay of treatment for early high-risk breast cancer patients, thus, uncovering mechanisms of chemoresistance is a crucial necessity.

To elucidate resistance mechanisms, the ideal way is to perform longitudinal studies, looking at matched pre- and post-treated tumors. The neoadjuvant (NAT) pre-operative setting provides a unique opportunity to study the effects of chemotherapy on real life tumors by the inherent existence of both pre-treatment diagnostic biopsy and post-treatment surgery specimen [13]. NAT are widely used for the treatment of high risk breast cancer patients and provide predictive and long-term prognostic information by evaluating tumor chemo-sensitivity [14]. The quest for biomarkers predicting response or for molecular targets avoiding resistance instigated many studies that inspected post-NAT residual tumors. While initial studies profiled one-time point primary tumor sample [15] or residual post-NAT tumors [16], later studies compared pre- and post-treatment tumors [17–21], resulting in the discovery of treatment-altered genes and pathways. Yet, most studies were based on statistical differential expression and could not provide a personalized view of resistance. More recent studies, inspected time-course changes following various neo-adjuvant treatment modalities, using matched serial sampling of three time points: pre-NAT, short-term and long-term post-treatment [6,13,22–25].

Nevertheless, the mechanisms underlying non-genetic resistance and the extent of phenotypic diversity between patients following treatment remains elusive. Transcriptional reprogramming in response to chemotherapy can be mediated by a stochastic mechanism that drives non-genetic heterogeneity [26]. Recent *in vitro* studies suggest that phenotypic resistance to therapy is achieved through drug-tolerant persister cancer cells, that their fraction is pre-determined in a non-stochastic manner [27,28]. It is unclear yet whether the non-genetic resistance involves active reprograming of resistant tumors or it was transcriptionally primed before treatment. To investigate this in clinically relevant samples, we collected a cohort of neo-adjuvant breast cancer patients and profiled their transcriptome before and after treatment, compared to the adjacent normal epithelium of the same patient. Thereby, we inspected drug-induced modulations relative to the deregulation state at the primary tumor. To understand the personal route to resistance we applied a patient-oriented pattern analysis algorithm for dealing with longitudinal datasets. We previously developed this pattern analysis approach to investigate miRNA expression modulations following recurrence [29], and further applied it to a longitudinal proteomic NAT dataset of breast cancer patients, revealing the involvement of proline biosynthesis in resistance [30].

The analysis of our longitudinal dataset depicted important principles in the adaptive non-genetic mechanism of resistance. The analysis yielded patient-specific chemoresistome maps, highlighting rewiring of simultaneous multiple pathways, albeit in a unique fashion to each patient. Personal chemoresistome mapping can provide insightful mechanistic understanding of therapy resistance with future clinical implication for rational design of treatment plans.

## Results

### Longitudinal dataset of breast cancer patients receiving neoadjuvant chemotherapy

Matched archived samples (pre-treatment tumor, post-treatment tumor and adjacent normal breast epithelia) were collected from a cohort of breast cancer patients that underwent neoadjuvant chemotherapy (Figure 1A). Tumor samples were collected from various subtypes and stages (full description of clinico-pathological characteristics are summarized in Table 1). Chemotherapy treatment included a standard-of-care sequential doxorubicin and cyclophosphamide followed by paclitaxel, and additional Herceptin for HER2+ patients. Post-treatment samples were scored for response by both Miller-Payne (MP) score [31] and RCB grade [32]. Whole transcriptome mRNAseq dataset was generated, using our optimized mRNAseq methodology, for archived FFPE samples [33]. A flow diagram presents the cases included in the analysis (Figure S1).

**Figure 1:**
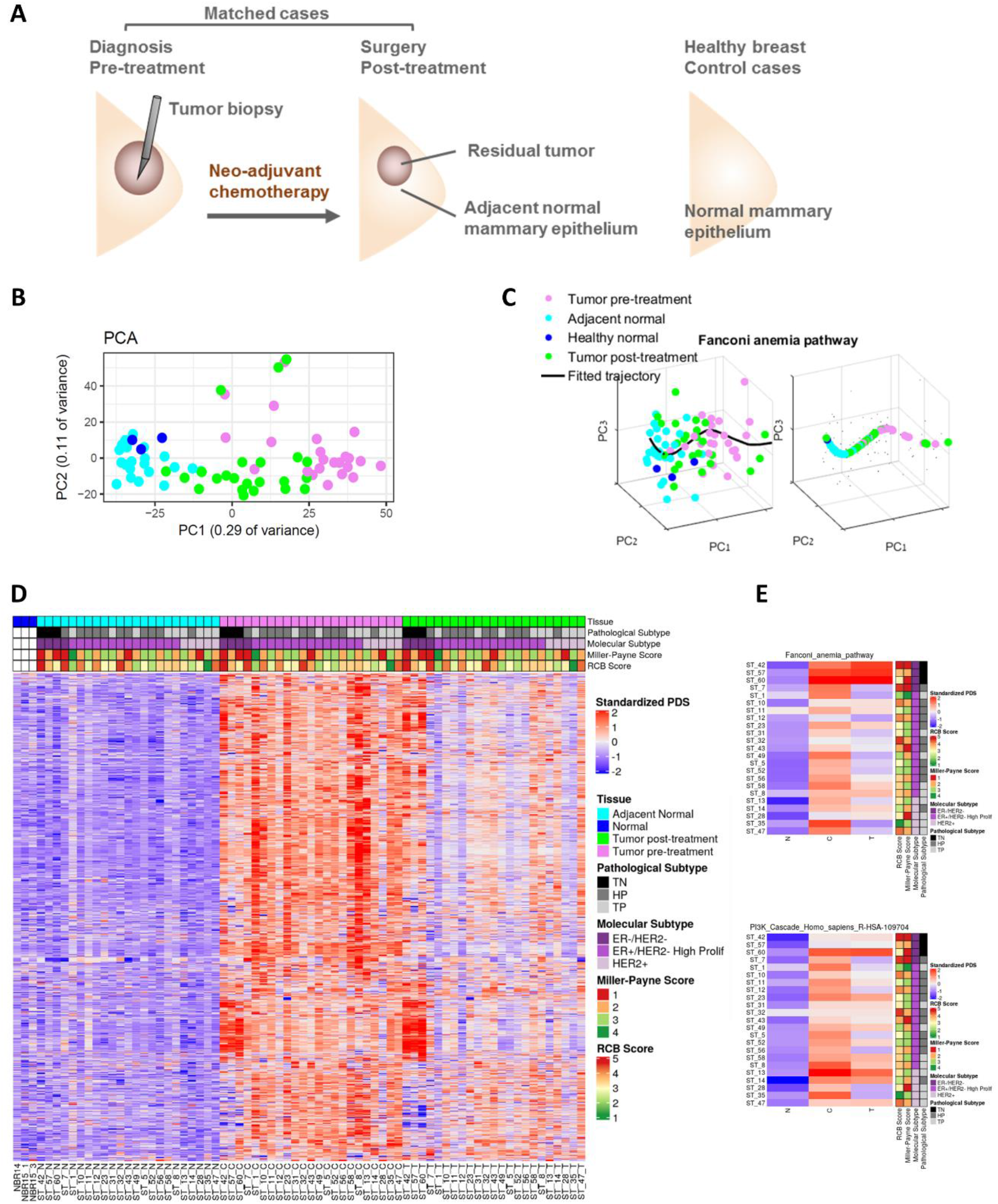
Longitudinal transcriptomics of matched samples from breast cancer patients receiving neoadjuvant chemotherapy. **A.** Archived samples from 29 breast cancer patients undergoing neo-adjuvant therapy were collected. From each patient we analyzed matched pre-treatment, post-treatment and adjacent normal breast epithelia. In addition, normal breast samples from 3 healthy women were included. **B.** Principal components analysis of the 1000 most variable genes from the whole transcriptome mRNAseq data. Gene expression clustered according to sample type. Adjacent normal breast samples were similar to normal mammary from healthy women. **C.** An example of calculating pathway deregulation score (PDS) using the Pathifier algorithm for a selected pathway from the KEGG dataset. The principal curve (black) is going through the cloud of points representing the various samples. The samples (colored according to their type) are projected onto the curve. **D.** Pathway Deregulation Scores (PDS) were calculated for each sample per pathway using the Pathifier algorithm. Each raw represents a different pathway (KEGG, Reactome and GO), for which a PDS relative to all normal samples is calculated. Each column represents a different sample in individual patient. Notably, healthy normal breast tissue (NBR) is very similar to the adjacent normal tissues (N) as shown by their overall low PDS (blue). Tumor samples pre-treatment show the most deregulated scores, as shown by their overall red levels. Tumor samples post-treatment are variable. E. Temporal modulations in deregulation scores per-patient for two representative pathways.

**Table 1:**
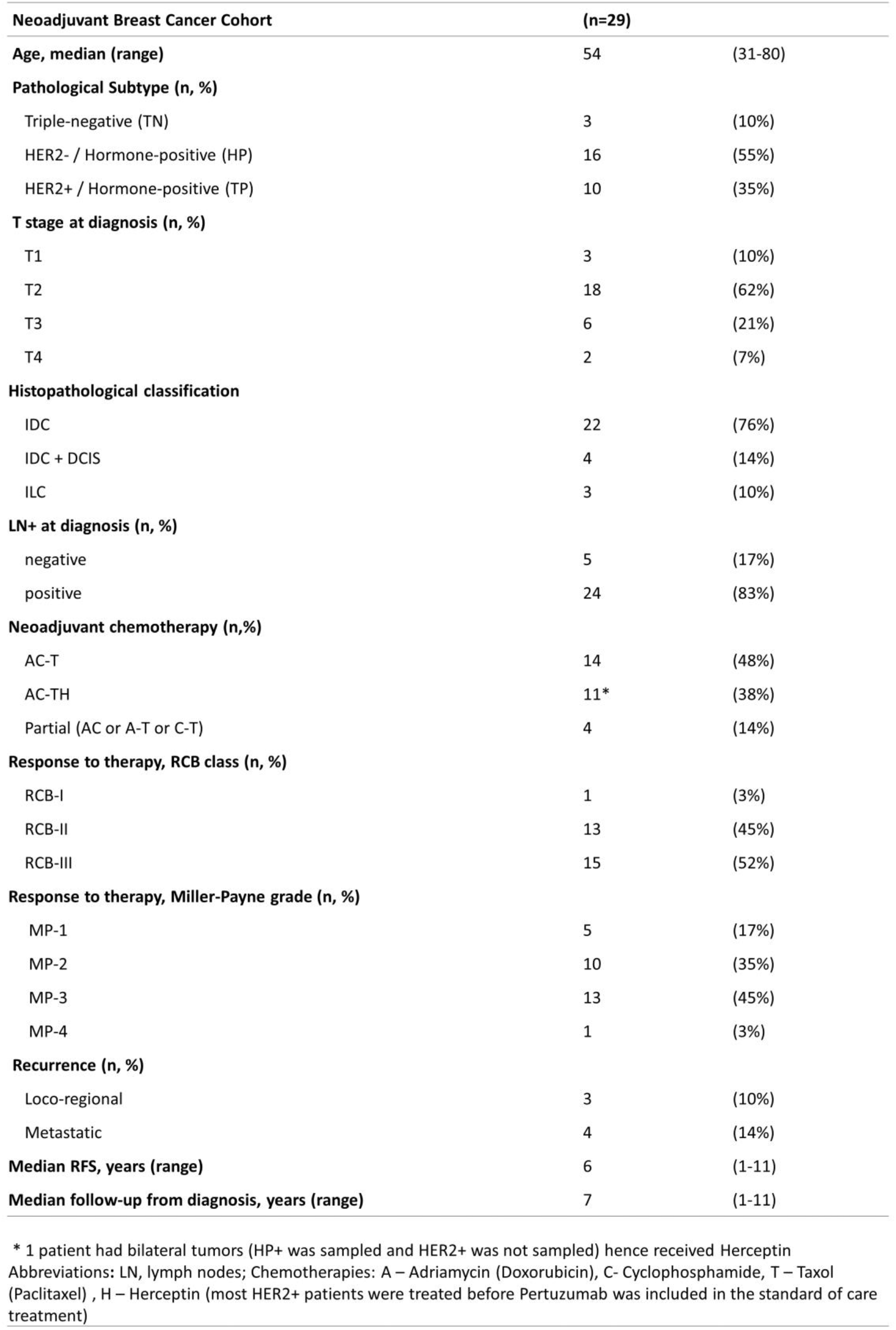
clinico-pathological parameters for the entire cohort

### Different sample types cluster together showing distinct regulation states

Expression-based Principal Component Analysis (PCA) of all samples demonstrated clustering of the samples according to sample type (pre- or post-treatment tumors and normal epithelium) (Figure 1B). Post-treatment samples were positioned between the adjacent normal tissues and the cancerous pre-treatment tissues, possibly reflecting the range of response to treatment. Notably, the healthy normal breasts were clustered with the adjacent normal samples of cancer patients (Figure 1B).

The overall modulations occurring through the course of disease and therapy were estimated by calculating pathway deregulation score (PDS) for the various samples, relative to all normal samples using Pathifier [34,35]. This algorithm calculates a PDS for each sample, based on the distance from the projection of the cloud of points on a principal curve (representative pathway exemplified in Figure 1C). PDS was calculated for a combined list of pathways (GO_biological process, KEGG and Reactome). A bird’s-eye view of the deregulation scores is shown as a heatmap of all pathways across all samples ordered by Molecular Subtype (Figure 1D). Notably, adjacent normal tissues were very similar to healthy normal breast tissues, indicating that these post-treatment normal samples largely represent normal breast epithelium. Pre-treatment tumors exhibited the most deregulated scores, whereas post-treatment tumors were variable. The overall map of pathway deregulation suggests that dynamic changes in specific pathways vary between patients. To examine this pathway-specific divergence in dynamics, we plotted PDS of matched samples for each patient (Figure 1E and Figure S3). We observed that while in some patients pathways returned to normal scores after treatment, in other patients pathways remained at the deregulated state. For example, the Fanconi anemia pathway is highly deregulated in the TNBC patients and remains deregulated also post-treatment, but in patients ST-7 and ST-1, PDS return to normal levels post-treatment. The patient-specific modulations in PDS suggests that a longitudinal analysis of the data would facilitate a deeper understanding of the adaptive processes during chemotherapy. To understand whether the variability among patients in the same pathway is attributed to either specific genes or diverse genes, we proceeded to analysis of temporal modulations at the gene level per-patient.

### Longitudinal pattern classification to identify genes associated with response

Inspecting the temporal gene modifications, we found that the dynamics of gene expression is predominantly patient-specific. Typically, genes showed divergent expression dynamics that were either response-related or subtype-related (Figure 2A and Figure S4). Only few genes showed patient-wide patterns (Supp. Table 2) (Figure S5A), with significantly higher expression levels in adjacent normal, also observed in the TCGA datasets, verifying that this difference is not due to treatment effects (Figure S5B).

**Figure 2:**
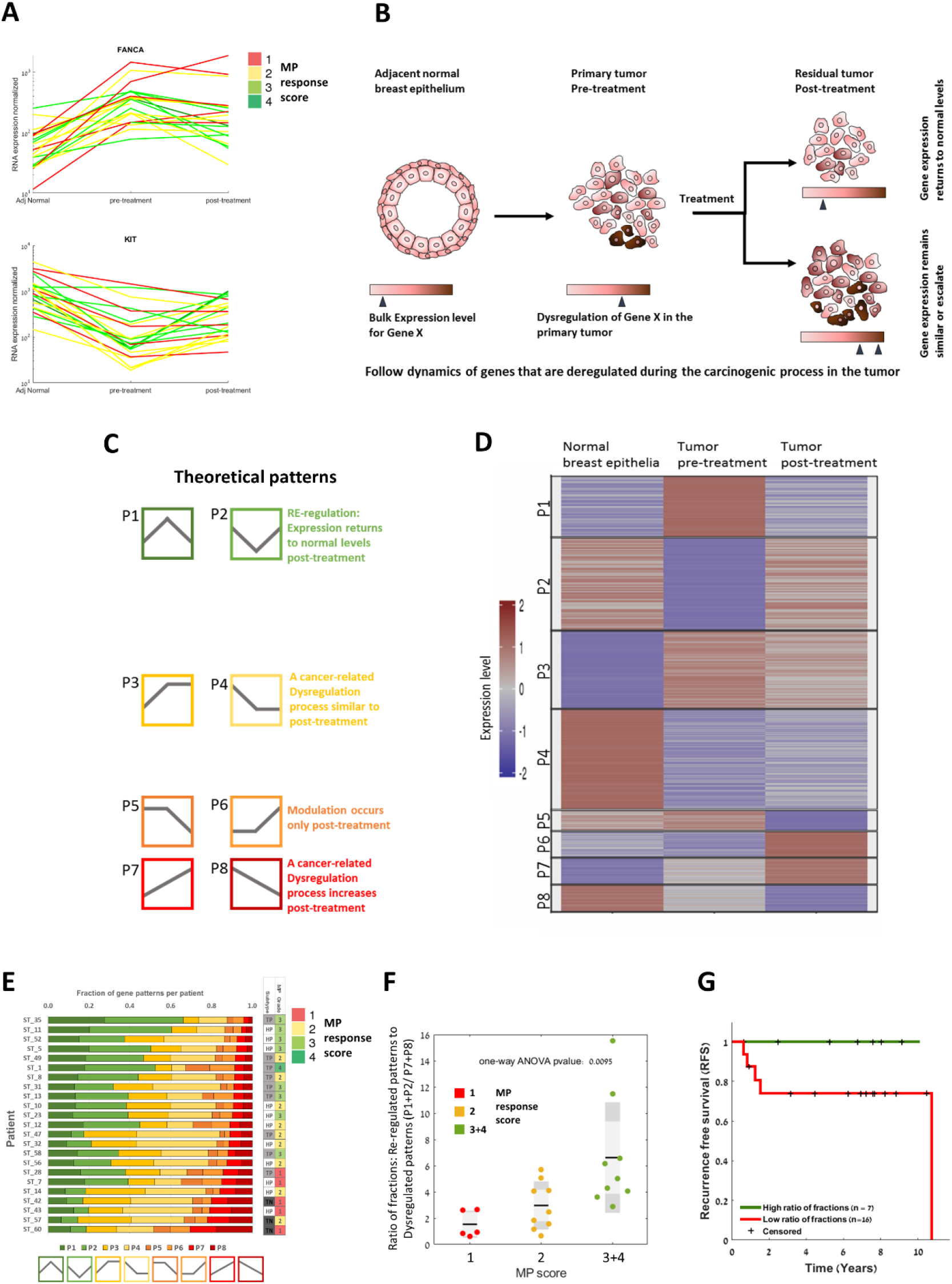
Longitudinal pattern classification to identify genes associated with response/resistance. A. Most genes showed divergent dynamics of expression between time points that were typically response-related. B. A scheme presenting the hypothesis that divergent dynamics in bulk expression can pinpoint genes that are associated with resistance to treatment. C. Differentially expressed genes were classified into eight possible theoretical patterns of modulations. The various patterns and their suggested scenarios are presented in different colors. D. Heatmap of all gene-per-patient triplets after classification by correlating the Variance Stabilizing Transformation values to the pre-defined theoretical patterns. Each row corresponds to one gene in a certain patient. The same gene is plotted for each patient, according to its classified pattern in an individual patient. E. Fractions of genes in the various patterns for each patient. Patients are ordered by the percentage of genes with patterns P7 and P8, which represent increased dysregulation post-treatment relative to the normal levels. F. Ratio of “re-regulated patterns” Pl and P2 to “dysregulated patterns” P7 and P8, (P1+P2/ P7+P8), was calculated for each patient. This ratio was significantly associated with MP response score (one-way ANOVA, p-value = 0.0095). G. Kaplan-Meier curve of recurrence free survival for the entire cohort by their ratio of fractions: (P1+P2/ P7+P8), cutoff = 5, Log rank p-value 0.09.

These few genes typically show highest co-expression of gene-pairs among patients in single-time points datasets (TCGA and METABRIC, Supp. Table 3 and Figure S5C), indicating on their robust regulation.

For most genes, the temporal patterns, rather than the absolute expression values, were correlated with treatment response score. For example, the absolute pre-treatment expression levels of *FANCA* or *KIT* (Figure 2A and Figure S4) did not separate between the various pathological MP response groups. Rather, the dynamic expression pattern differentiated between response groups (Figure 2A and Figure S4). This divergence in dynamics led us to hypothesize that classifying genes by their expression pattern can identify resistance-related genes, as depicted in a scheme in Figure 2B. We thus performed dynamic pattern analysis for the differentially expressed genes, similar to our previous studies [29,30], exploiting the strength of the longitudinal dataset. We defined eight possible theoretical patterns (P1-P8) of gene expression modulations through the matched triplet of normal-tumor-treatment stages (Figure 2C). Each pattern represents a different scenario of events through tumor progression and treatment stages:

1. Increased (P1) or decreased (P2) expression levels from normal to pre-treated sample and return to normal levels post-treatment. This may represent re-regulation of genes, responding to treatment.
2. Increased (P3) or decreased (P4) expression levels from normal to pre-treated sample, which remain similar post-treatment, possibly representing resistance state.
3. Similar expression between normal and tumor samples, which decrease (P5), or increase (P6) only post-treatment, that may be inferred as treatment effects.
4. Constant increase (P7) or constant decrease (P8) in expression levels from normal through pre-treated sample to post-treatment sample. This may represent a worsening in the cancerous process.

Genes-per-patient trajectories were classified into the various patterns by correlating the Variance Stabilizing Transformation (VST) values (Examples of theoretical patterns and assigned genes in Figure S6). A heatmap presenting the distribution across patterns of all gene-per-patient trajectories is shown in Figure 2D. The same gene is plotted for all patients, according to its patient-specific classified pattern. To validate the classified pattern directions, we compared the fold changes between tumor-normal pairs to fold changes observed in the METABRIC datasets (fold change of 1.5 and FDR <= 0.05) [8] and found an agreement in the direction of upregulation or downregulation in tumor vs. normal tissue.

### Longitudinal gene pattern fractions are associated with response

To examine the association between expression patterns and response, we plotted the fraction of genes assigned to each pattern per-patient (Figure 2E). Patients with better response score (higher MP value) showed higher fractions of genes with patterns P1 and P2, in which expression returns to normal levels.

Poor responders showed higher fractions of P3 or P4 and mainly P7 or P8, suggesting an increase in cancer-related processes post-treatment. Notably, cellularity of cancer cells in the sample was not associated with the fraction of patterns. We found that the ratio between the fractions of genes with patterns P1 and P2 relative to P7 and P8 (P1+P2/ P7+P8), was significantly associated with MP response score (ANOVA test, p-value = 0.0095) (Figure 2F). This fraction of patterns was also associated with recurrence free survival, although not significant due to the small cohort size (Figure 2G). The MP score for this cohort was also associated with recurrence free survival (Figure S7, ns), emphasizing its clinical validity [31].

To understand how the various patterns are distributed among patients, we clustered the matrix of number of patients per pattern for all differentially expressed genes (~6900 genes). We observed that frequently, genes were classified into combinations of two main patterns across patients (Figure 3A). Genes that tend to be upregulated in cancer were classified either as P1 in some patients or P3 (and few as P7) in other patients. Genes that tend to be downregulated in cancer, were classified into either P2 or P4 (and few as P8). Only few genes were classified into patterns P5/P6, scarcely shared among patients. We speculated that this dichotomous distribution among patients could differentiate between good responders and poor responders (Figure 3A). We thus calculated the correlation between gene pattern and MP response score for genes with dichotomous patterns (Wilcoxon Rank sum test), and identified 253 significant genes associated with response/resistance to treatment (Figure 3B) (Suppl. Table 4).

**Figure 3:**
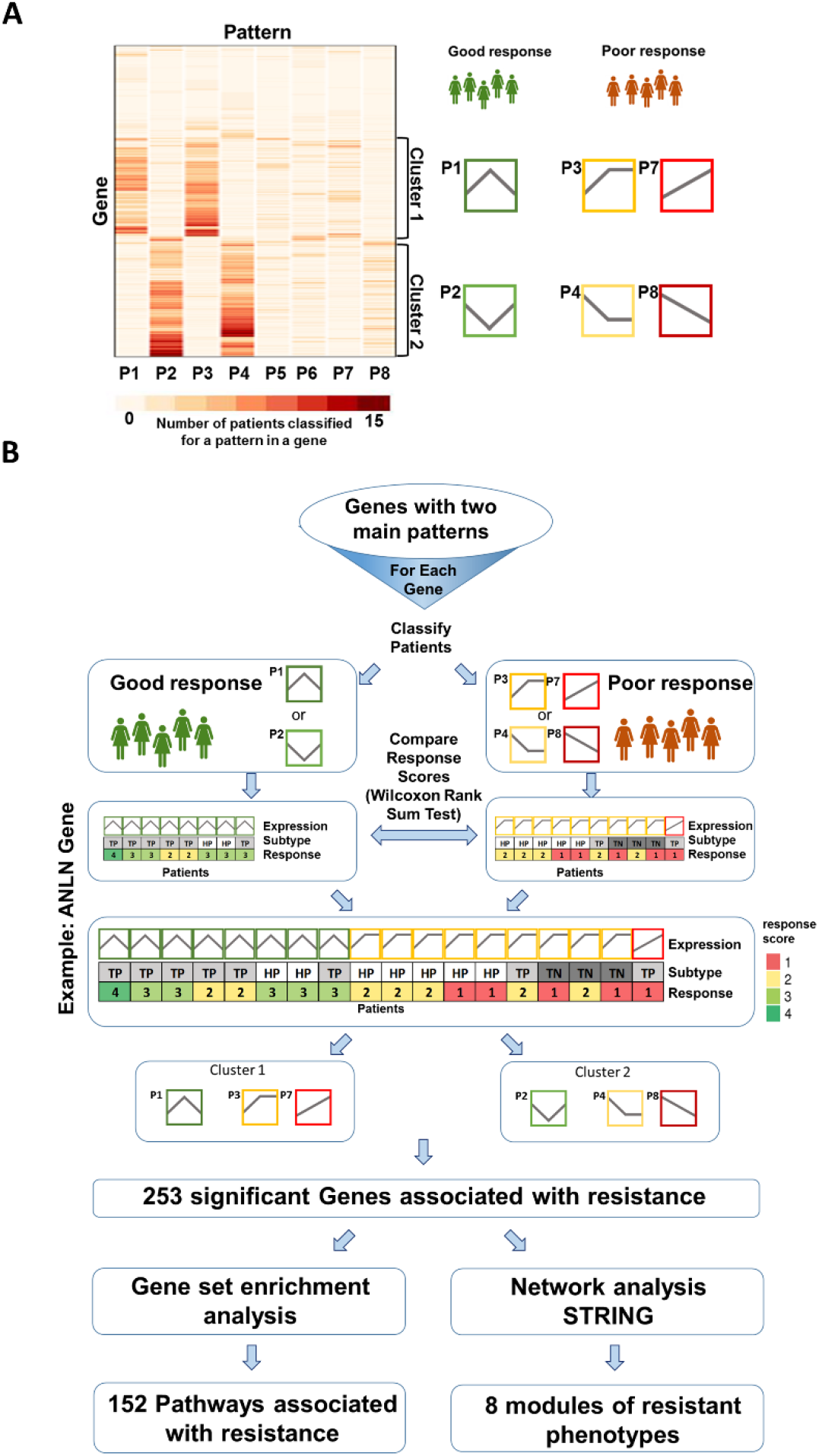
Gene patterns analysis identifies genes and pathways associated with resistance. A. Differentially expressed genes typically exhibit dichotomous assignment to patterns (two main patterns), namely, the same gene is either assigned to two different patterns (for example, Pl or P3). B. Flow chart of the gene pattern analysis algorithm. For genes with dichotomous patterns, we compared the MP pathological response score of the patients (using the Wilcoxon Rank sum test). We identified 253 genes that their pattern is significantly correlated with response. These genes were further analyzed by enrichment analysis, identifying 152 pathways associated with resistance. Further functional analysis of these genes included network and gene set enrichment analyses..

### Resistance genes converge to a finite number of chemo-resistance functional modules

To explore the functional context of the resistant genes, we performed gene-set enrichment analysis of these genes (EnrichR) [36] using the combined datasets (GO_biological process KEGG and Reactome), and identified 152 pathways significantly associated with resistance (q value < 0.05) (see Suppl. Table 5). Pathways were related to cell cycle, mitosis, DNA replication and DNA repair as well as to MAPK and PI3K cascade, angiogenesis, lipid transport and extracellular matrix organization.

To further understand the functional connectivity between the identified 253 resistance-related genes, we performed STRING network analysis. We found that the genes converge to a finite number of resistance modules (Figure 4A). Notably, three modules were related to the mechanism of action of the administered chemotherapies (Taxane, cyclophosphamide and Doxorubicin)[37]. Taxanes act by disrupting microtubule function thereby inhibiting mitosis. The divergent dynamics of STMN1, previously shown to be associated with resistance to Taxanes [38], was significantly correlated with MP response (p-value 0.006) (Figure S4, top left). Cyclophosphamide exerts its cytotoxic effects mainly by cross-linking DNA strands. We identified five genes that were enriched in the DNA repair pathway (Reactome) (Suppl. Table 5), mainly in the Fanconi anemia complex (Figure 4A). Other emerged resistance modules were related to ECM remodeling, PI3K and AKT signaling, angiogenesis regulation, glucose transport, the ABCA2 family of the ABC transporters, associated with steroid transport, and lipid metabolism (Figure 4A).

**Figure 4:**
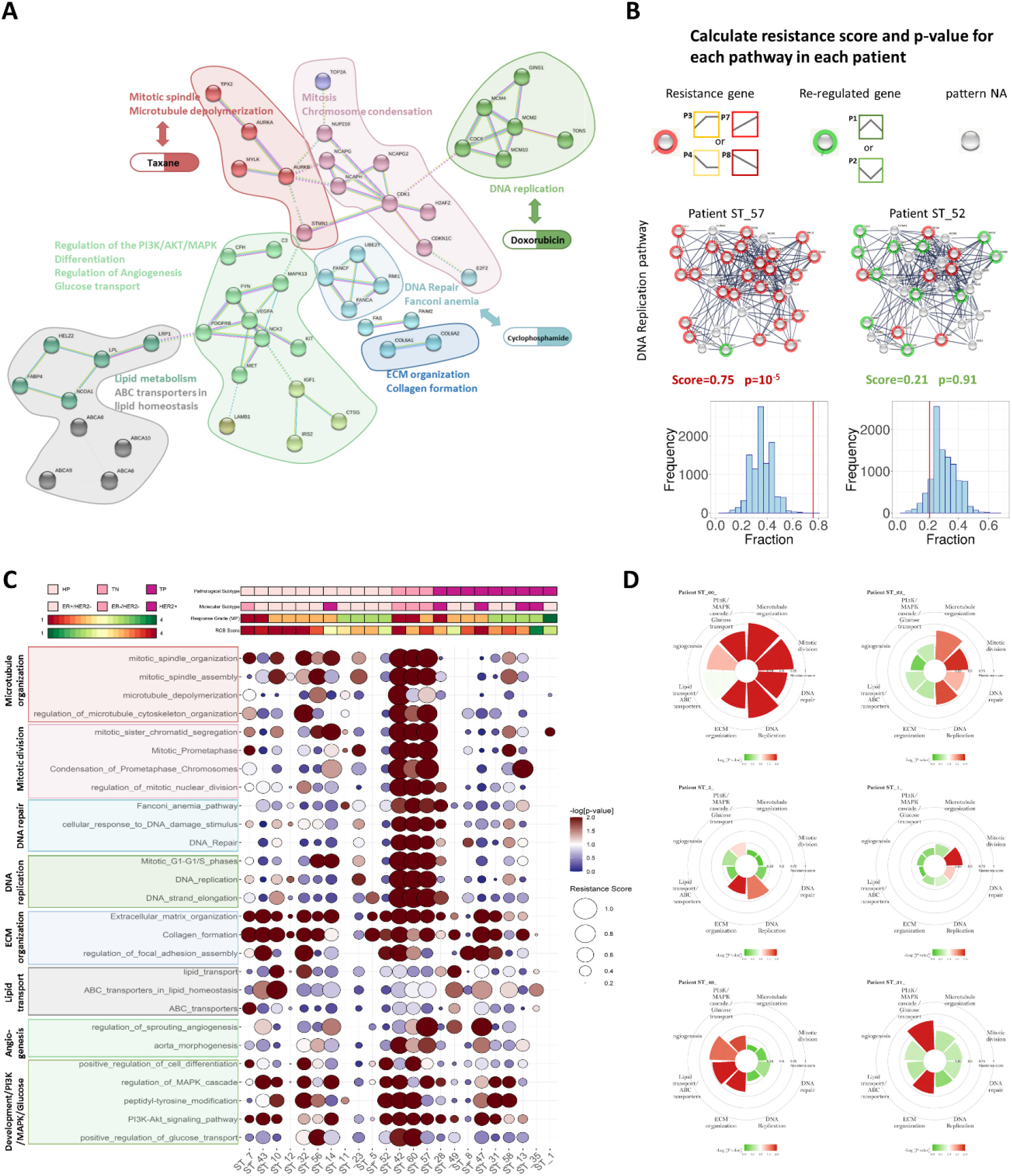
Functional modules and pathways associated with resistance are divergent across patients. A. Network analysis by STRING for all the genes associated with response demonstrates the convergence of the resistance genes into a confined number of resistance categories. Importantly, three categories are strongly associated with the mechanism of action of the administered chemotherapies. B, Enrichment analysis identified 152 resistance-related pathways. For each pathway in each patient, genes were assigned a mode of either resistance or re-regulation based on their classified pattern. We then calculated a resistance score: fraction of “resistant genes” in the pathway and a p-value for this score, calculated by bootstrapping a random set of genes in the same size of the pathway (sampled 10,000 times). An example for the calculated resistance score in two representative patients for the DNA replication pathway, presented as a network. Red dots: resistance genes, green dots: re-regulated genes, grey dots: genes with no assigned pattern. The p-value calculation is presented as a histogram of the distribution of random fractions in this patient. The red line denotes the resistance score {fraction of resistant genes in this pathway for this patient). C. Resistance score and p-value for 27 representative pathways enriched for resistance-related genes presented as a baloonplot. Each row represents a pathway and each column represents a patient. Circle size denotes the resistance score and color-code denotes the p-value of the resistance score. Clinical parameters for each patient are color-coded in the upper panel. The pathways are gathered according to their functional module, in the same color as in the STRING network in A. D. Personal chemoresistome maps for representative patients exemplifying the divergence in adaptive resistance for each patient. The maps represent the main resistance modules, calculated for the most significant pathways in each module. The resistance score is denoted by the bar height and the p-value is color-coded.

### Individual patients exhibit distinct dysregulation of diverse genes within pathways

To understand the divergence in dysregulated genes between the patients, we inspected all expressed genes in the 152 dysregulated pathways, for each individual patient. We defined genes with patterns P1/P2 as “re-regulated genes” and genes with patterns P3/P4/P7/P8 as “resistance genes”. We observed that the fraction and identity of resistance genes in the same pathway varied between patients. We therefore calculated for each patient a resistance score – fraction of resistance genes per pathway. The significance of this score was calculated by bootstrapping random sets of genes in the same size of the pathway in each patient (sampled 10,000 times). Calculation of resistance score for two patients is exemplified by a network map of a representative pathway (Reactome: DNA Replication), and the p-value is represented by histograms of the random fractions of resistance genes for this pathway (Figure 4B).

The resistance score and p-value per-patient for a curated list of 27 significant resistance-associated pathways is presented by a balloon plot, demonstrating the variability in dysregulated pathways between patients (Figure 4C and Suppl. Table 6). The pathways were categorized into the eight resistance modules identified in the network analysis and are colored according to the module (Figure 4A). Importantly, it is evident from the plot that in some patients multiple pathways account for resistance, while in other patients only few pathways exhibit high resistance scores. A clear difference between subtypes is evident in TNBC patients (ST_42, ST_57 and ST_60), exhibiting multiple resistance-related pathways typically associated with cell division, while in HR+ patients, resistance-related pathways vary and are specific to each patient. For example, patient ST-11 and patient ST-35 show significant resistance score in only few pathways, i.e., the Fanconi Anemia pathway in ST-11 and the ABC transporters pathway in ST-35 (figure 4C). In contrast, patients ST-10, ST-32 and ST-7 exhibit significant resistance scores in multiple pathways. This divergence in specific dysregulated pathways in response to the same chemotherapies pinpoint the individual emergence of adaptive resistance.

To highlight the distinct contribution of each module to the emergence of resistance in individual patients, we generated personal chemoresistome maps by circular bar-graphs (Figure 4D). The maps exemplify the co-existence of several resistance modules/mechanisms in each patient, while highlighting inter-patient variability by the impact of each module. Importantly, some modules exhibited patient-specific or subtype-specific occurrence, emphasizing the individual emergence of resistance. We observed that resistance modules related to cell cycle and cell division are abundant in TNBC and fast-dividing tumors, while other modules, such as ECM remodeling and PI3K/AKT regulation, were more characteristic of HR+ tumors. Nevertheless, cell proliferation modules were also evident in non-TNBC patients, but are frequently more sporadic and specific to a mechanism-of action of one drug.

### Variability in gene and pathway dysregulation reveals the principles in adaptive chemo-resistance

To further explore the inter-patient variability within dysregulated pathways, we zoomed into the various genes in each resistance-associated pathway. The inter-patient variability of the rewired genes is exemplified for two pathways by a gene network maps (Figure 5A) and by individual chemoresistome map (Figure 5B). For example, in the Fanconi anemia pathway, many genes exhibited resistance patterns in patient ST_42 (TNBC), while in patients ST_7 (HR+) and ST_8 (TP) the number of resistance genes varied (Figure 5A). Specifically, the genes in this pathway for patient ST_7 exhibit mostly re-regulated patterns, although all patients received Cyclophosphamide. The inter-variability between these patients was also evident in the microtubule depolymerization pathway. When comparing the resistance genes between patients we found that there are hubs of genes (Patterns P3, P4, P7 and P8), that recur in many patients (Figure S8). For example, in the Fanconi anemia pathway, RMI2 and FANCA exhibited resistance patterns in 13 patients (Figure S9A and S9B). Importantly, patient ST_52 did not receive paclitaxel and exhibited only re-regulated genes in the microtubule depolymerization pathway (Figure S9C), suggesting that resistance-related rewiring is induced by Taxane, that targets this pathway. However, the variability between patients is high and we observed only re-regulated genes in other patients (mostly HR+/HER2+) that received paclitaxel (Figure S9C).

**Figure 5:**
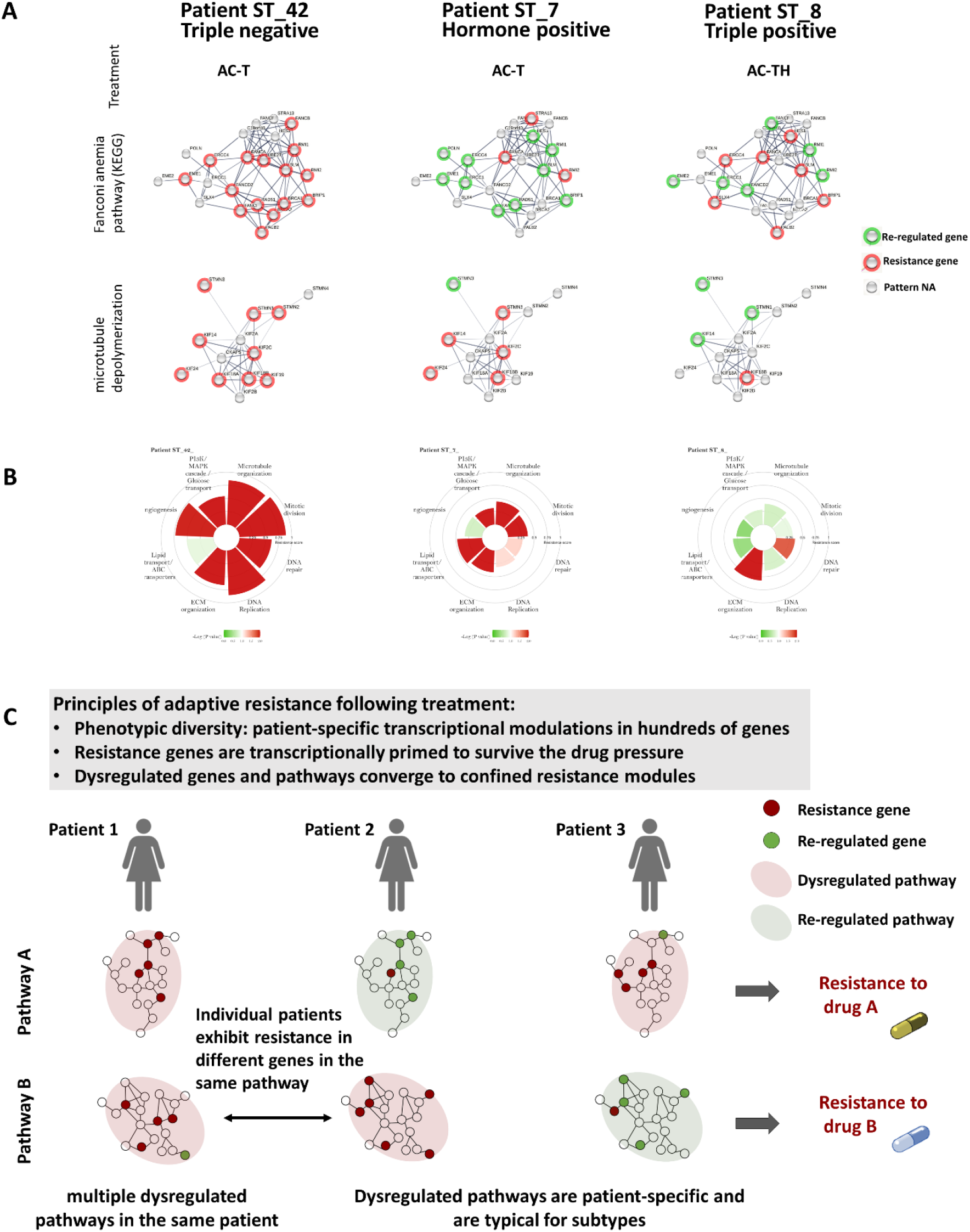
Variability in gene rewiring and pathway dysregulation reveals the principles of adaptive resistance following chemotherapy. **A**, An example for the variability in pathway dysregulation between patients. Network presentation of the expressed genes in two representative pathways, colored by the gene pattern analysis to classify genes into either re-regulated or resistance genes (see Figure 4B). While patient ST_42 (TNBC) exhibited multiple resistance genes in both pathways, patient ST_7 and patient ST_8 exhibited opposite resistance tendency for the two pathways. **B.** Personal chemoresistome maps for the corresponding patients in A, presenting the significant resistance modules in the patients. The resistance score is denoted by the bar height and the p-value is color-coded. C. Principles of adaptive resistance following chemotherapy. Hundreds of genes undergo rewiring following chemotherapy, in a patient-specific manner. The resistance-related genes are frequently transcriptionally primed in the primary tumor. The same pathway can undergo dysregulation by rewiring different genes. The rewired genes and dysregulated pathways converge into confined resistance modules, typically related to the mechanism of action of the administered drugs.

Altogether, the results of this study led us to suggest the principles of adaptive resistance post-chemotherapy (illustrated in Figure 5C). We propose that hundreds of genes undergo rewiring following chemotherapy. The rewired genes are mostly cancer-related genes that were already modulated during the carcinogenic process relative to the normal epithelium tissue, some of which return to normal levels while others maintain dysregulated levels. Although many genes and pathways exhibit dysregulation post-treatment, they converge to a finite number of confined resistance modules. Furthermore, rewired genes and dysregulated pathways are patient-specific, thus different genes in the same pathway can be rewired in specific patients.

## Discussion

The results of this study provide a wide perspective on the complexity of cancer cells adaption to chemotherapies. Inspecting patient-specific modulations between pre- and post-treatment tumor samples, relative to the normal epithelium of the same patient, revealed that rewiring of gene expression is a composite interplay, unique to each tumor. We postulate that multiple pathways, already dysregulated in the primary tumor, maintain a drug-tolerant state by activating many genes that safeguard their abnormal activity. Thereby, to survive the toxic activity of a drug, tumor cells either sustain the dysregulated state or intensify it, specifically bypassing the drug’s interference.

Our approach of longitudinal sampling, combined with pattern analysis, emphasized the importance of inspecting dynamic changes to reveal mechanisms of drug resistance. Most resistance genes could not be detected in a single time-point analysis, as their absolute levels in the primary tumor were indistinguishable between good and poor responders. For example, the TOP2A gene, specifically targeted by Doxorubicin, was previously evaluated as a marker for benefit from this drug [39], but inconsistent results impaired its predictive efficacy. Our data clearly indicates that TOP2A expression levels in the primary tumor are inconsistent with response, but its longitudinal expression pattern differentiates between good and poor responders (Figure S4). This may explain why heterogeneous tumor phenotypes challenge the identification of molecular markers and suggests that patient-specific dynamic changes are crucial to target the resistance genes and pathways.

A key advantage of our dataset is the inclusion of matched adjacent normal tissue. In previous longitudinal studies, the search for differentially expressed genes before and after treatment missed non-modified genes that sustain a dysregulated state, as in Pattern P3. On the other hand, genes that are frequently modified post-treatment but are actually re-regulated (patterns P1/P2), can be misinterpreted as markers for chemo-resistance (exemplified in a scheme, Figure S10). For example, CYR61 was previously reported to be upregulated post-NAT [22], and was suggested as a chemo-resistance marker. In our dataset, its levels are indeed upregulated post-NAT, but to similar levels as in adjacent normal tissue, thus inferred as a re-regulated gene (Figure S10A). In contrast, upregulation of FN1 post-NAT was previously associated with poor response [40] and mainly exhibited resistance patterns in our dataset (Figure S10B).

The fact that most resistance genes were identified in patterns P3/P7 or P4/P8, modulated in the carcinogenic state relative to the normal epithelium, suggests that tumors resist treatment by maintaining dysregulated pathways. Signaling pathways that are crucial for cancer cells survival sustain their activity or shift to higher gears to persist under chemotherapy extreme conditions. Our observations suggest that persistent subpopulations possess primed transcriptional profiles, relative to the normal state, that convey gene expression bias towards survival. This notion is in accordance with recent *in vitro* studies of drug-tolerant cells, showing that phenotypic resistance is determined a priori in a non-stochastic manner in sub-populations of persister cells [27,28]. Another study of single-cell sequencing of TNBC patients in response to chemotherapy supports this concept, showing clonal selection of pre-existing genotypes, as well as transcriptional reprogramming of resistant signatures in the persistent clones [6]. Similar to our results, transcriptional programs converged to a few resistance pathways and were mostly acquired after treatment. Our data confirms the concept that resistant phenotypes were already acquired in the primary tumor, relative to matched normal tissue, and their dysregulation had either persisted or intensified.

Dynamic changes in response to chemotherapy may occur as a result of genetic selection or by massive transcriptional adaptation. The kinetics of transcriptional adaptation are considerably faster than genetic evolution, and it is assumed that transient non-genetic mechanisms will eventually be hardwired through long-term epigenetic or genetic mechanisms [2]. *In vitro* pattern analysis of time-course omics during treatment revealed immediate massive transcriptional adaptation followed by concordant epigenetic alterations, suggested to stabilize the resistant phenotype [41]. Notably, our study measures modulation in the bulk expression levels that can represent either transcriptional selection of cells expressing a gene or a transcriptional adaptation by gene rewiring. Either way, the net levels point toward the dysregulation or re-regulation state of the tumor. As shown in our study and in previous studies [42], gene rewiring converges to a confined set of drug-resistant phenotypes. However, while most studies focus on genes and pathways that are shared between patients, we emphasize the phenotypic heterogeneity between patients and the divergent routes of adaptive resistance.

The emergence of key signaling pathways in our dataset, directly related to the mechanism of action of the chemotherapies, resembles the mechanism of resistance to kinase inhibitors, by which persistent activation of the drug target bypasses its inhibitory effect [1]. Nevertheless, the results also indicate an acquisition of an additional resistance mechanisms, such as angiogenesis, ECM-related, PI3K/AKT signaling and lipid metabolism alterations. While in TNBC tumors proliferation pathways are mainly upregulated (P1), in HR+ tumors differentiation pathways are downregulated (P2). In TNBC tumors dominant resistance pathways were related to cell cycle, mitotic spindle, Hedgehog pathway, DNA replication and DNA repair, indicative of these highly proliferative tumors. In HR+ tumors we mainly observed resistance patterns in PI3K/Akt pathways, regulation of MAPK cascade, ECM-related pathways, lipid transport and ABC transporters pathways. Indeed, Non-genomic activity of estrogens has been suggested to induce chemoresistance, activating PI3K/AKT and modifying DNA damage response [43]. Resistance patterns emerged in specific subsets of ABC transporters, the ABCA2 family, involved in steroid transport that may regulate breast cancer proliferation and survival, as was suggested in a prostate cancer study [46].

Our findings may have several future clinical implications. Overcoming drug-tolerant persister cell populations in the clinical setting is challenging, given that there is no one driver mutation to target. Our data suggests that while many resistant phenotypes are emerging in parallel, re-regulated pathways disclose their sensitivity, enabling to rationally select the next therapeutic plan. Residual tumors after NAT have higher incidence rate for recurrence [14]. These residual tumors harbor distinct molecular profiles that disclose their acquired resistance on one hand and their vulnerabilities (i.e., re-regulated genes upon treatment) on the other hand. Therefore, profiling the chemoresistome immediately after surgery, may guide the adjuvant treatment plan of targeted therapy to prevent recurrent or metastatic disease. Alternatively, future developments such as cell-free RNA or epigenetic profiling of tumor DNA can lead to chemoresistome profiling during the initial cycles of the neoadjuvant treatment. Moreover, recent advances allow for profiling of primary tumors ex-vivo adjacent to diagnosis [47]. For cost-effectiveness it may be useful to profile the chemoresistome and not all the transcriptome. Furthermore, such defined resistome maps can be calculated for immunoresistome or other targeted therapies.

The results of our study are limited by the small cohort size and diverged subtypes. The findings and suggested principles should be further validated in larger cohorts as well as in other treatment regimens. Though the cohort size was too small for subgroup analysis, the combined subtypes analysis was advantageous for detecting patient-specific modulations. Many resistance-related modulations that were subtype-specific, mainly in TNBC, were found in few patients from other subtypes. Although we enriched our samples for epithelial cell by macro-dissection, confounding factors that may impair the inference of our analysis are related to stroma-associated changes after treatment, including overrepresentation of collagen and immune cells.

In summary, elucidation of principles governing mechanisms of adaptive resistance to chemotherapy has been facilitated by our unique longitudinal dataset coupled with pattern analysis. A complex and heterogeneous phenotypic adaptation depicted from this study appears as a major obstacle for successful anti-cancer treatment. However, the insightful conclusions may accelerate optimization of patient-specific treatment plan according to the observed adaptive states.

## Methods

### Cohort assembly

The cohort consisted of 29 breast cancer patients who underwent neoadjuvant therapy, and did not achieve complete response, resulting in partial response and available residual tumor. Only patients with partial response were included, as we aimed to focus on dynamic changes between the primary tumor and the residual tumor. Clinicopathological information for the cohort is summarized in Table 1 and detailed in supp. Table S1. For each patient we collected triplets of pre-treatment tumor (diagnostic biopsy), post-treatment residual tumor (sampled at surgery, approximately 6-8 months after the pre-treatment biopsy) and adjacent normal epithelium (sampled from the post-treatment surgery specimen). Samples were collected as Formalin-Fixed Paraffin-embedded (FFPE) tissues from the Sheba Medical Center pathology archive. Additional normal breast epithelium samples were collected from breast reduction specimens of six healthy individuals. Overall, a total of 97 samples were included in this study. Two independent pathologists inspected the slides to define tumor cellularity and scored response by Miller-Payne (MP) score [31] and by RCB (Residual cancer burden class) [32]. Recurrence free survival according to Miller-Payne scores was performed by Kaplan-Meier analysis, calculating log-rank p-value. The study was approved by the Institutional Review Board (IRB) (approval #8736-11).

### mRNAseq library preparation

FFPE tumor samples were macro-dissected to enrich for tumor cellularity. The 5 microns sections were deparaffinized at 90°C for 5 min. Total RNA was extracted using nucleic acid isolation kit (AllPrep® Qiagen) according to protocol instructions. RNA concentrations were determined by Qubit™ fluorometer (Thermofisher scientific). Whole transcriptome profiling of the archived samples was accomplished by a reliable and cost-effective mRNAseq procedure that we previously optimized for FFPE samples [33]. Briefly, mRNA-seq libraries were constructed by Truseq RNA Sample preparation kit v2 (illumina). Libraries size was evaluated by Tapestation (Agilent). Primer dimers were eliminated using 1x Agencourt RNAClean XP Beads. Sequencing libraries were constructed with TruSeq SBS Kit and multiplexed to run 6-10 samples per lane with a read length of 60 bp single-end run in Illumina HiSeq V4 instrument. The median sequencing depth was ~27 million reads per sample.

### Sequence alignment and mapping

Illumina adapters and PolyA/T stretches were trimmed using cutadapt [48]. Reads shorter than 30 bp were discarded. The reads were mapped to the human genome (GRCh38) using STAR v2.4.2a [49]. Overall, mapping of reads to the genome was high, with an average of 87% reads being mapped and 61% of the uniquely mapped reads being counted. Median uniquely mapped reads were 10.7 million reads. Counting proceeded over genes annotated in Ensembl release 88, using HTSeq-count [50]. Out of the 97 samples, four samples from normal epithelium failed the library preparation steps due to low RNA quantity. Additional six samples failed QC after mapping and three samples were excluded due to low tumor cellularity. Overall, 84 samples from 29 patients were included in the general analysis. Finally, 23 patients had complete triplet samples that passed all data QC and were used for pattern analysis (see Figure S1 for flow chart of samples). Pipeline was constructed using Snakemake [51]. Data analysis was mostly done by R language [52] using the specified packages.

### mRNAseq data evaluation and molecular subtype calculation

The dynamic range of the mRNAseq data was correlated to pathological scoring values of immunohistochemistry staining for: ER, PR, HER2 and KI67 when available (Figure S2A,B). Percentage of Ki67 positive cells were assessed blinded to the expression data using the automated Virtuoso image analysis algorithm (Ventana). The dynamic range of the mRNAseq data at highest pathological scores for the receptors was much higher than the immunohistochemistry staining levels (Figure S2C).

To evaluate the purity of the tumor samples from the mRNAseq counts, we used the ESTIMATE algorithm [53] and compared it to tumor purity, estimated by two independent pathologists after the macro-dissection. Overall, there was a good agreement between the pathological evaluation and the purity calculated by the ESTIMATE algorithm.

Molecular subtype was calculated from the mRNAseq data by the Genefu package [54] using the SCMOD2 subtyping algorithms [55] (detailed in supp. TableS1).

### Differential Expression Analysis

To analyze the differential expression between matched sample types (normal - pre-treatment - post-treatment) we utilized DESeq2 package [56] with the betaPrior, cooksCutoff and independentFiltering parameters set to False. Raw P values were adjusted for multiple testing using the procedure of Benjamini and Hochberg. To estimate the number of differentially expressed genes between each pair of sample types, the following contrasts were defined: post-treatment tumor vs. normal, pre-treatment tumor vs. normal and post-treatment vs. pre-treatment. The genes were filtered to keep genes with an absolute fold change above 1.5, adjusted p-value below 0.05, and a count of at least 100 in at least one of the samples. About 6900 genes passed this threshold.

### Pattern classification

To analyze the modification of genes across the three time points (normal, pre-treatment and post-treatment) we defined all possible eight theoretical patterns (Figure 2C) as was previously published [29].

To assign a pattern for each gene we correlated the changes in expression values, calculated by DESeq2 Variance Stabilizing Transformation (VST), to the theoretical Patterns. VST calculates sample geometric means, estimates dispersions for each gene, fits a mean-dispersion trend, and then transforms the data to stabilize the variance across the mean. Each gene was assigned to a theoretical pattern (according to the maximal correlation). Genes were considered as classified into one main pattern if they were classified to a single pattern in at least nine patients and if the proportion of this most frequent pattern was at least 20% higher than other patterns. Genes were considered as classified into two main patterns if the gene was classified to each of the two patterns in at least 30% of the patients and each of the two patterns was observed at least in five patients, based on the distribution of the patterns frequencies.

### Calculate overall fraction of patterns in each patient

For each gene classified with two main patterns (dichotomous behavior), we calculated the association of patterns with MP response score. Patterns P1 and P2 were considered as “good response”, while patterns P3, P4, P7 and P8 were considered as “poor response”. The fraction of genes classified to each pattern in each patient was correlated to pathological response rate. Ratio of P1+P2 (expression returns to normal levels) to P7+P8 (cancerous expression state worsens after treatment), was calculated for each patient and correlated with pathological response score.

### Defining genes associated with the response

In order to detect genes that are associated with response, we focused on genes with two main patterns. For each gene, we compared the Miller-Payne score of the patients with “good response” patterns (P1 or P2) versus scores of patients with “poor response” patterns (P3, P7 or P4, P8). Wilcoxon ranks sum test was used to calculate the significance of the pattern association with response. The test was calculated for each gene cluster separately: P1 vs P3/P7 and P2 vs P4/P8 (Figure 3B).

### Pathifier algorithm

We used the Pathifier algorithm [34,35] to calculate pathways deregulation scores (PDS) for each patient’s sample for a combined list of pathways databases: GO_biological process (2018), KEGG (2019) and Reactome (2016), all downloaded from EnricheR website [36]. Briefly, Pathifier analyzes each pathway, and assigns to each sample i and pathway P a score that estimates the extent to which the behavior of pathway P deviates from normal samples. To determine PDS, we used the normalized counts of all 97 samples. For each pathway, PDS is defined as the distance of each sample from its projection on the calculated principal curve of all samples to the projection of the normal samples (Figure 1C). Normal samples were defined as both the healthy normal and the adjacent normal samples.

### Pathway enrichment analysis and resistance score calculations

Pathway enrichment analysis for genes that were significantly associated with response (Wilcoxon rank-sum test, p-value <0.05) was calculated using ClusterProfiler package [57], using pathways from the following databases: GO_biological process (2018), KEGG (2019) and Reactome (2016) (downloaded from EnrichR database [36] website).

For each enriched pathway, we calculated resistance score per patient and a p-value. The resistance score is defined as the fraction of “resistance genes” (genes with patterns P3, P4, P7 or P8) in the pathway. Only genes that were significantly differentially expressed (see Differential Expression Analysis section) were considered. The p-value for this score, was calculated by bootstrapping random sets of genes of the same size of the pathway (sampled 10,000 times). For each patient in each pathway, the p-value was calculated by dividing the number of random scores, which were greater or equal to the observed score, by the number of permutations.

For a simple representation of resistance modules in each patient, we generated personal chemoresistome maps. The enriched pathways were reduced to a list of 27 pathways, accounting for significance and functional redundancy. These pathways were divided into functional modules. Circular barplots were plotted for each patient, representing the selected resistance modules. The resistance score and p-value in each module in the circular barplot is represented by its most significant pathway.

### STRING analysis

To identify the functional network of the resistance genes (253 genes, p-value by Wilcoxon rank test < 0.05) we ran STRING analysis (version 11.5) [58]. Interaction score was set to the highest confidence (0.9). K-means clustering (defined for 11 clusters) was used to identify functional categories. Functional categories were defined by enrichment of each cluster by reactome and GO biological process. For visualization of resistance genes in each pathway, all genes in the pathway were uploaded to the STRING’s payload mechanism and a color-code was specified for a halo around the STRING nodes.

## Acknowledgements

Acknowledgments

This research was supported by a grant from the Israel Ministry of Science (# 3-11175). This study makes use of data generated by the Molecular Taxonomy of Breast Cancer International Consortium (Curtis et al, Nature 2012). The Consortium Data Access Committee approved application for data access. G.F. is the Incumbent of the David and Stacey Cynamon Research fellow Chair in Genetics and Personalized Medicine. We thank Jonathan Barlev and Barak Markus for helpful discussions regarding data analysis.

## Authors’ Contributions

M. Dadiani designed the study concept, supervised the study, achieved funding, performed bioinformatics and statistical analysis, created visualization and wrote the manuscript. G. Friedlander performed bioinformatics and statistical analysis, created visualization and edited the manuscript, G. Perry collected the biopsies and the clinical data, processed the samples and edited he manuscript. N. Balint-Lahat performed pathological evaluation. S. Gilad designed the protocols and supervised the transcriptomics sequencing. D. Morzaev-Sulzbach quantified the immunohistochemistry staining. N. Bossel Ben-Moshe was involved in initial bioinformatics analysis and QC. A. Shenoy collected the samples and edited the manuscript. A. Pavlovsky processed the samples. I. Barshack supervised the pathological evaluation. T. Geiger was involved in study conceptualization, achieved funding and edited the manuscript. B. Kaufman (deceased) was involved in study conceptualization and achieved funding. EN Gal-Yam was involved in data interpretation, provided resources and edited the manuscript.

## Data availability

The RNA-seq data have been deposited in the database of Gene Expression Omnibus (GEO) and are accessible through GEO Series accession number GSE217624 (https://www.ncbi.nlm.nih.gov/geo/query/acc.cgi?acc=GSE217624).

Supplementary tables are available at the figshare data repository: https://figshare.com/s/cca0b5864aaaf7606f19

