## Supplementary material for "Chemoresistome Mapping in Individual Breast Cancer Patients Unravels Diversity in Dynamic Transcriptional Adaptation": supp. figures

Dadiani M et al.

### Extended data


- **Figure S1:Flow chart of patients and samples used in the analysis**
- **Figure S2: Correlation between expression levels and scores of pathological markers**
- **Figure S3: Temporal modulations in deregulation scores per-patient for representative pathways**
- **Figure S4:Diverged temporal expression patterns associated with resistance**
- **Figure S5: Genes with shared pattern dynamics across patients**
- **Figure S6: Pattern classification by correlating genes to Theoretical Patterns**
- **Figure S7: Kaplan-Meier curve of recurrence free survival by MP response score**
- **Figure S8: Hubs of resistant genes in selected dysregulated pathways**
- **Figure S9: Heat maps presenting the modes of resistance/reregulation in two representative dysregulated pathways**
- **Figure S10: Resolving resistance by pattern analysis of matched three time points.**

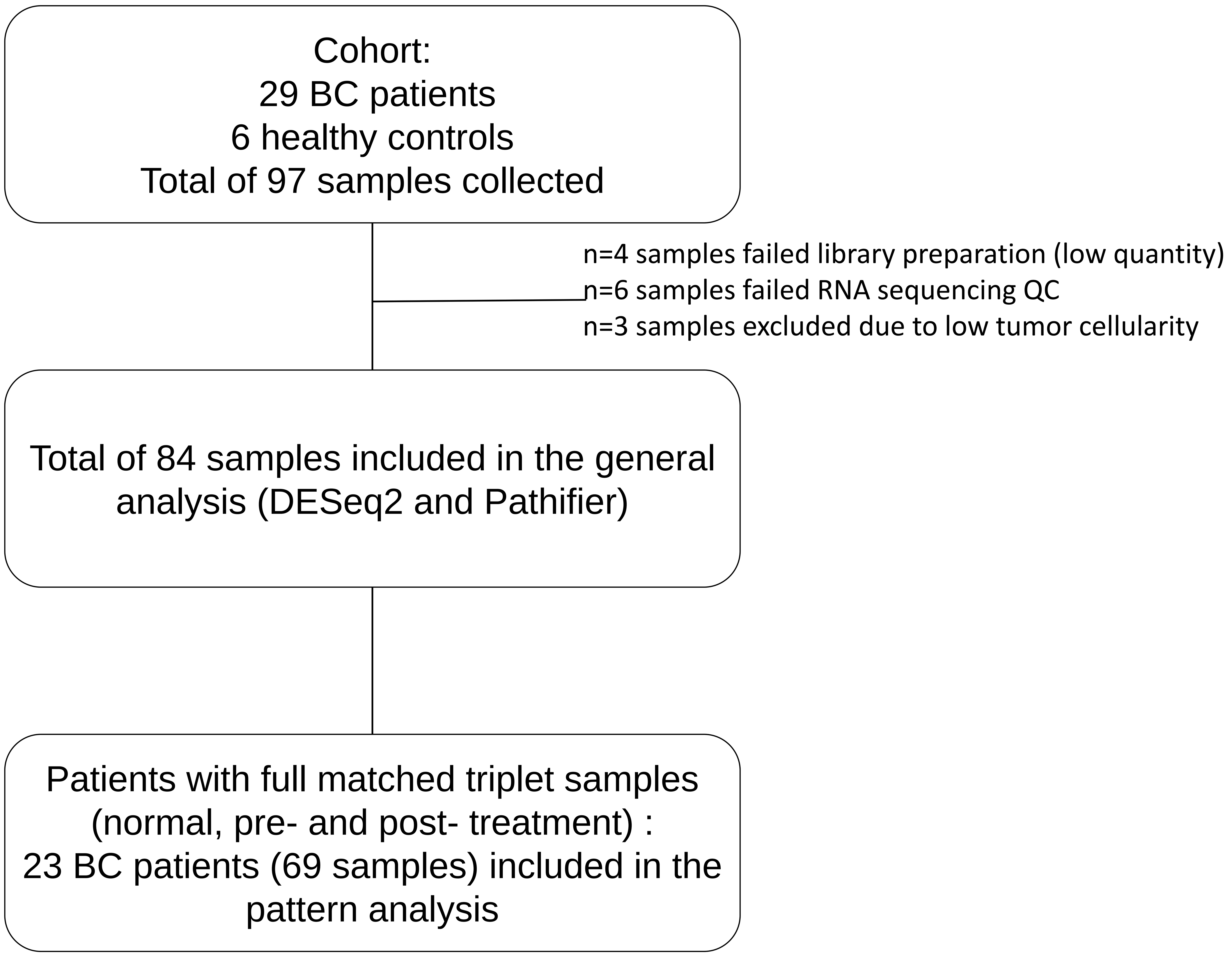


**Figure S1: Flow chart of patients and samples used in the analysis**

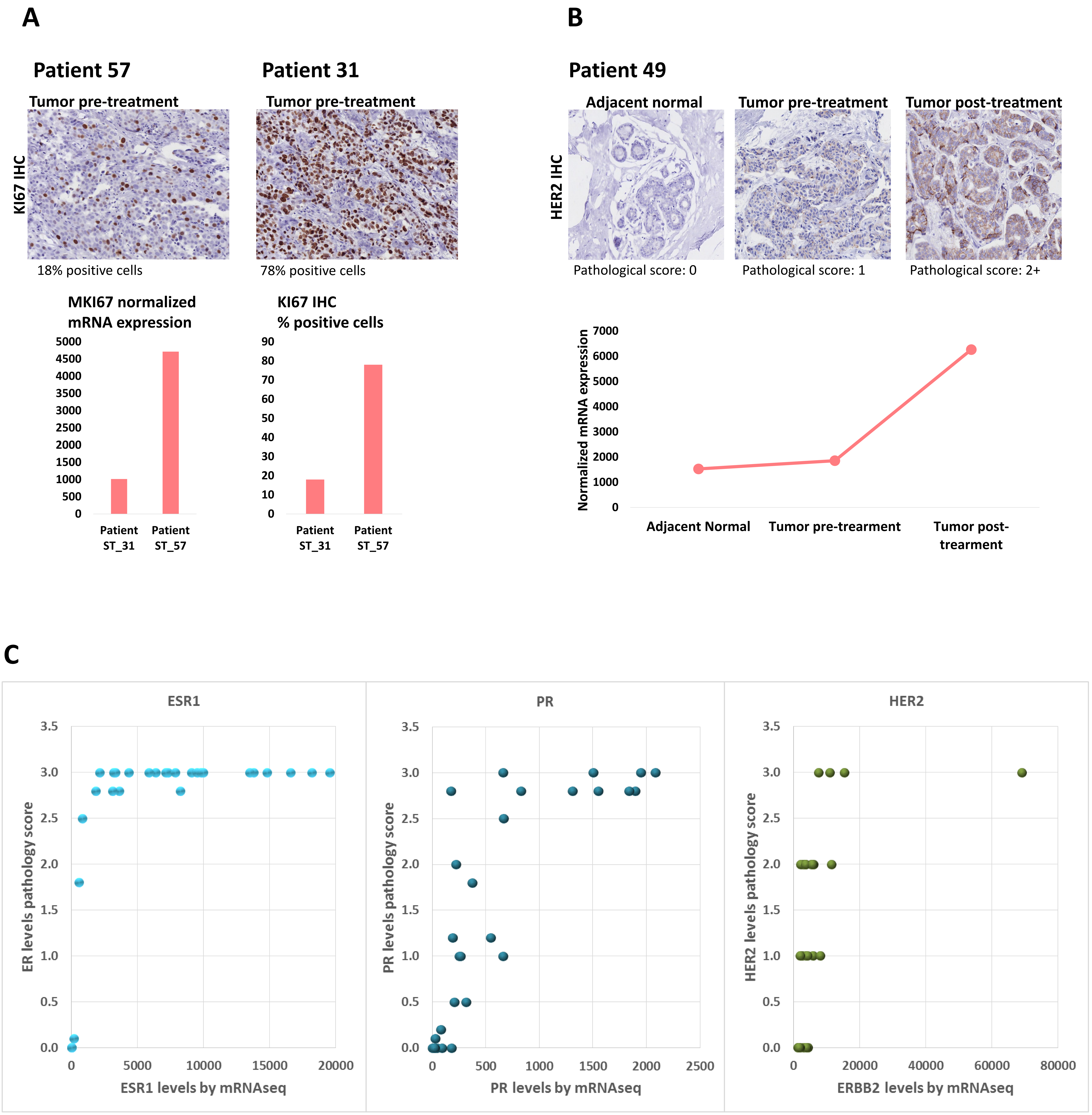


**Figure S2. Correlation between expression levels and scores of pathological markers. A.** MKI67 expression (normalized counts) of patients 57 and 31 are shown in red and green lines, respectively, for the three samples (adjacent normal, pre-treatment and post-treatment; x-axis). Percentage of Ki67 positive cells were assessed blinded to the expression data using the automated Virtuoso image analysis algorithm. B. ERBB2 expression levels (normalized counts and matched HER2 immunohistochemistry staining and its corresponding pathological score). C. The pathology score for each pathological marker is plotted against the normalized counts (RNA Seq) for the proteins ESR1 (left), PR (middle) and HER2 (right).

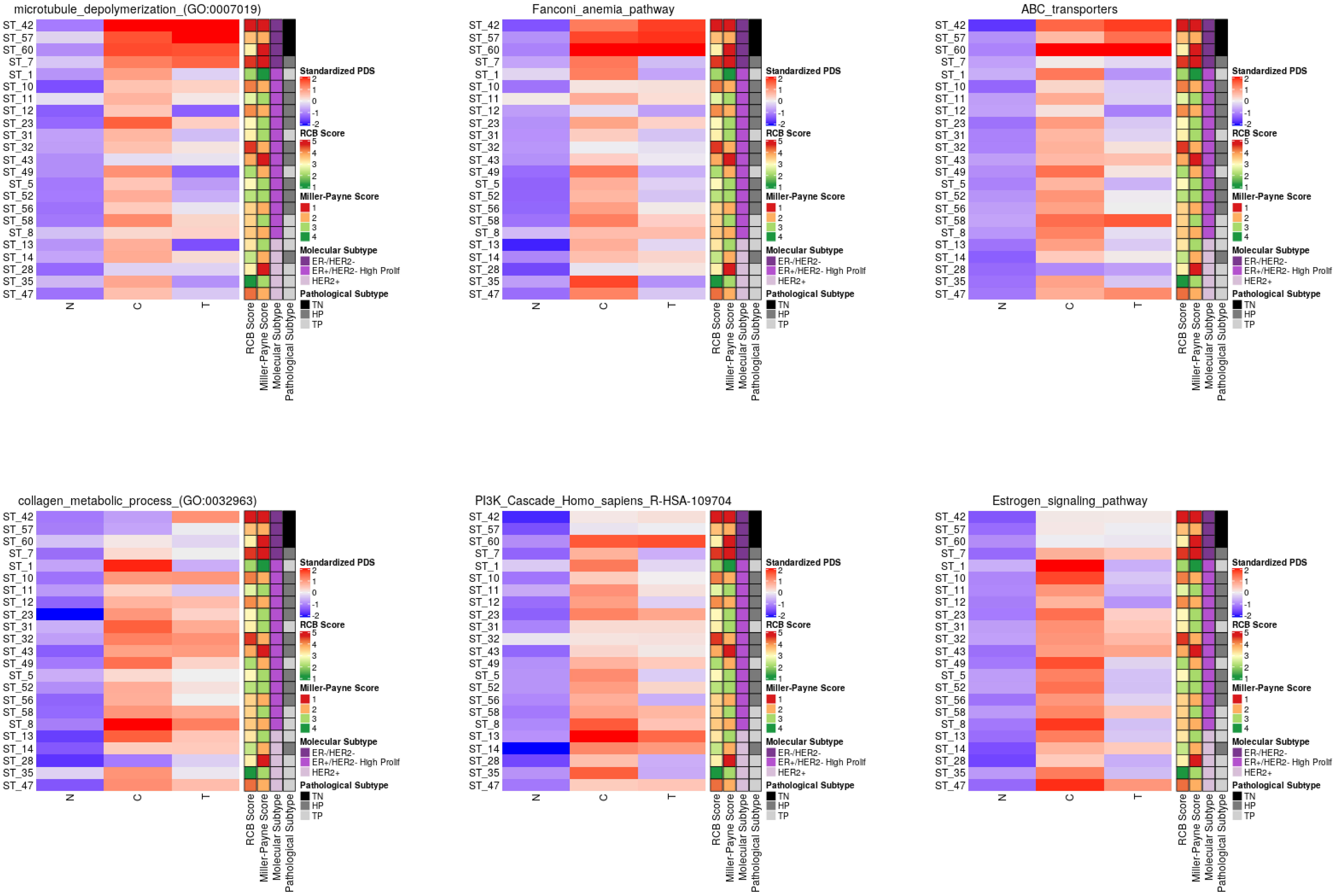


**Figure S3: Temporal modulations in deregulation scores per-patient for representative pathways.** Longitudinal representation of the pathway deregulation scores (PDS) calculated by Pathifier [43]. A heatmap of the PDS values, standardized to have for each patient zero mean and unit standard deviation. Each row relates to a patient. The columns relate to the tissue type (N, C or T).

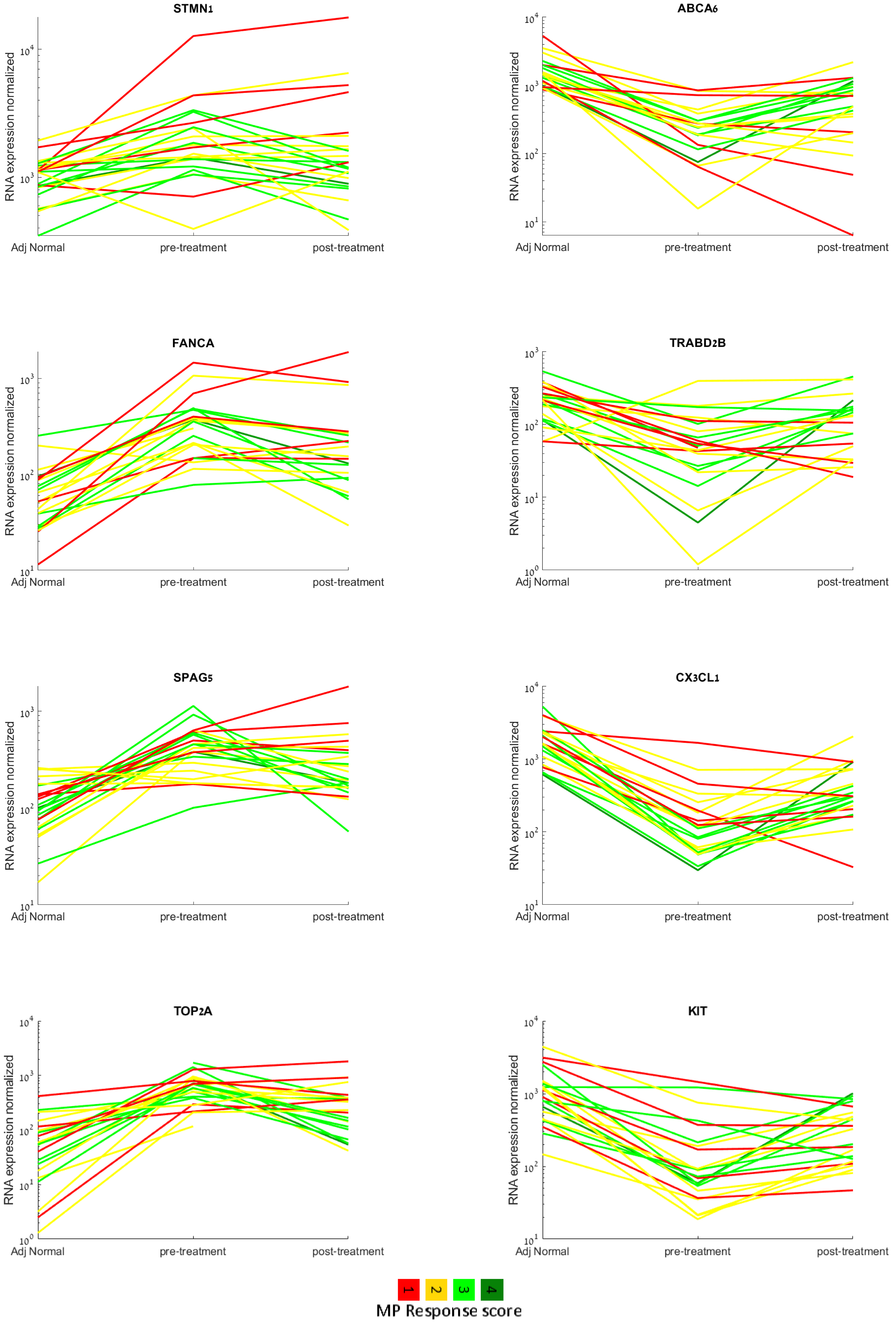


**Figure S4: Divergent temporal expression patterns associated with resistance**. Bimodal expression dynamics of representative genes that were significantly associated with pathological MP response score (by Wilcoxon Rank sum test).

**A. Patient-wide pattern dynamics** **B. Comparing tumor to adjacent normal expression levels (TCGA)** **C. Co-expression across patients (METABRIC) relative to FOS**

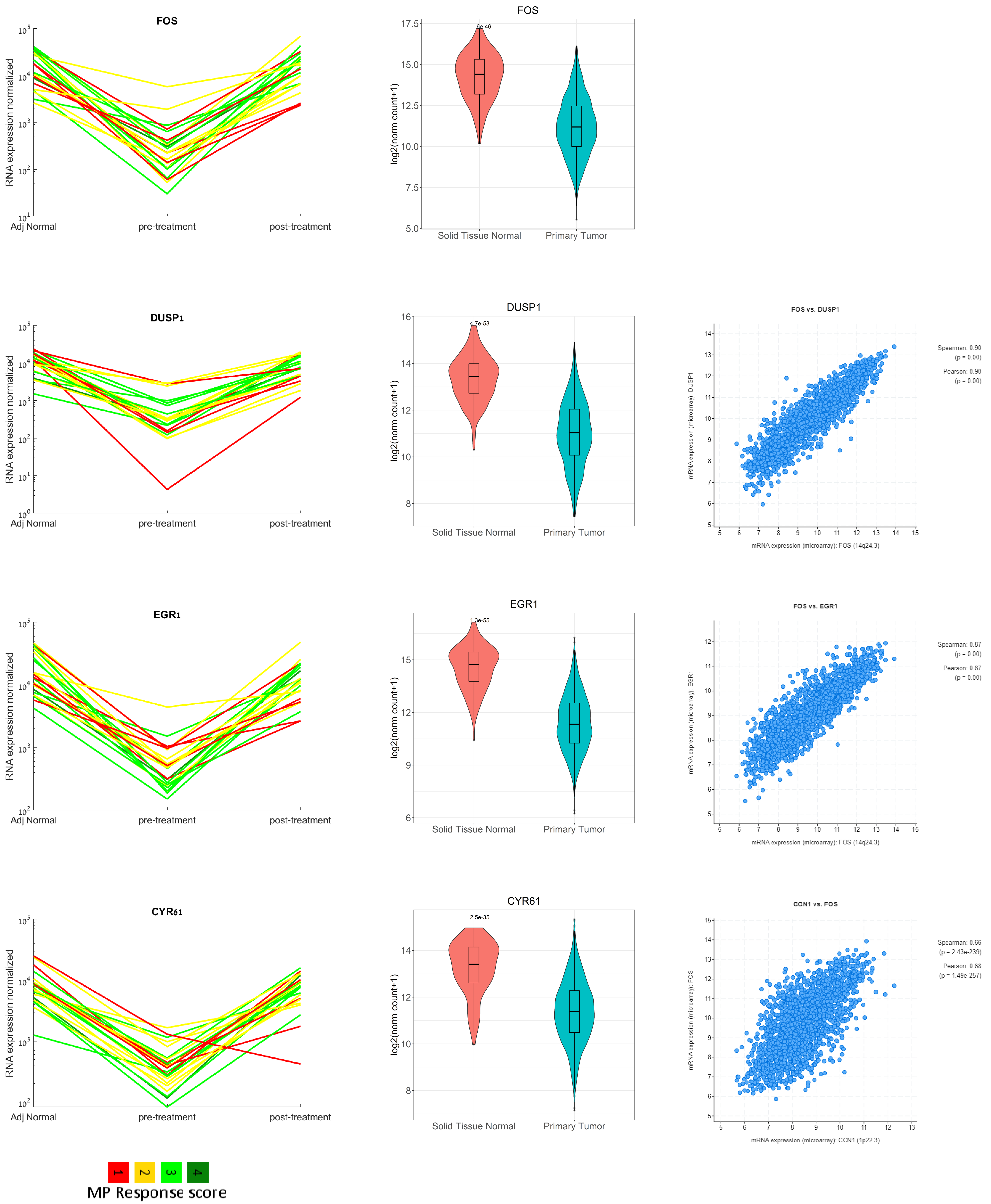


**Figure S5: Genes with shared pattern dynamics across patients.** A. Few genes exhibited the same pattern across all patients, independent of response or subtype. Each line denotes one patient, colored by the MP response score. B. Comparing expression values (log2(norm_counts+1) of tumor samples (n=1101) and adjacent normal samples (n=139). TCGA of breast cancer data downloaded from the XENA server. Adjusted p values (FDR corrected) are presented for each gene. C. These genes were found to have the most significant co-expression correlations across patients in large datasets (both TCGA and METABRIC). co-expression plots are derived from METABRIC dataset and plotted via cbioportal.

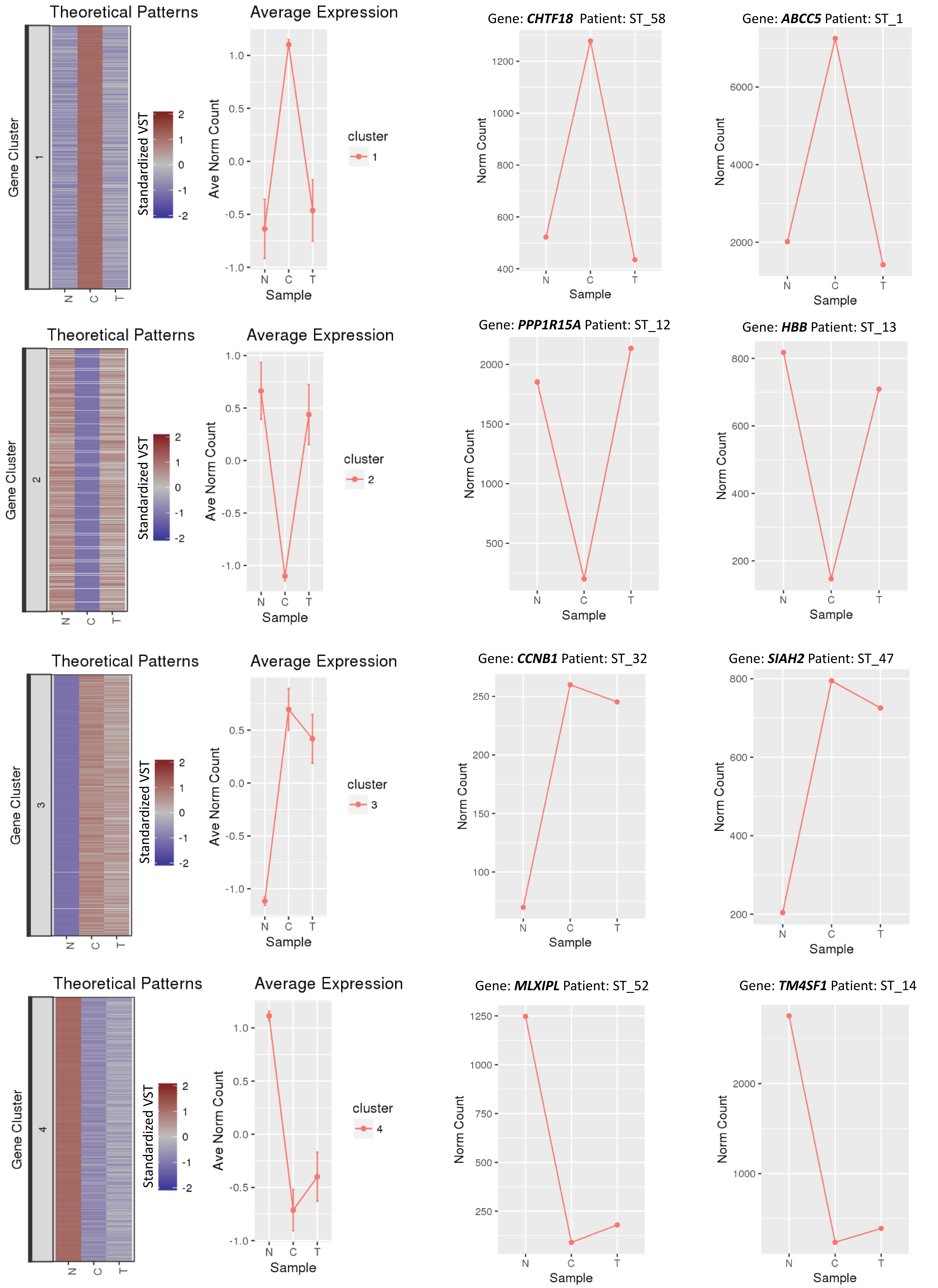


**Figure S6. Pattern classification by correlating genes to Theoretical Patterns .** The variance stabilized transformed (VST) expression values were correlated to the eight theoretical patterns**.** Each gene was assigned to a theoretical pattern (according to the maximal correlation). The left heatmap presents the expression for all the gene-patient profiles that were assigned to the pattern in the three tissue types (N, C and T). Shown are VST expression values, standardized to have for each gene zero mean and unit standard deviation. The expression profile is accompanied by a colored bar indicating the standardized values. The line plot shows the average expression of the standardized VST values. The error bars represent the standard deviation. On the right, shown are line plots of the Normalized counts of a gene in a single patient, in the three sample types (N, T and C) for two representative genes that were assigned to each of the patterns (P1-P4).

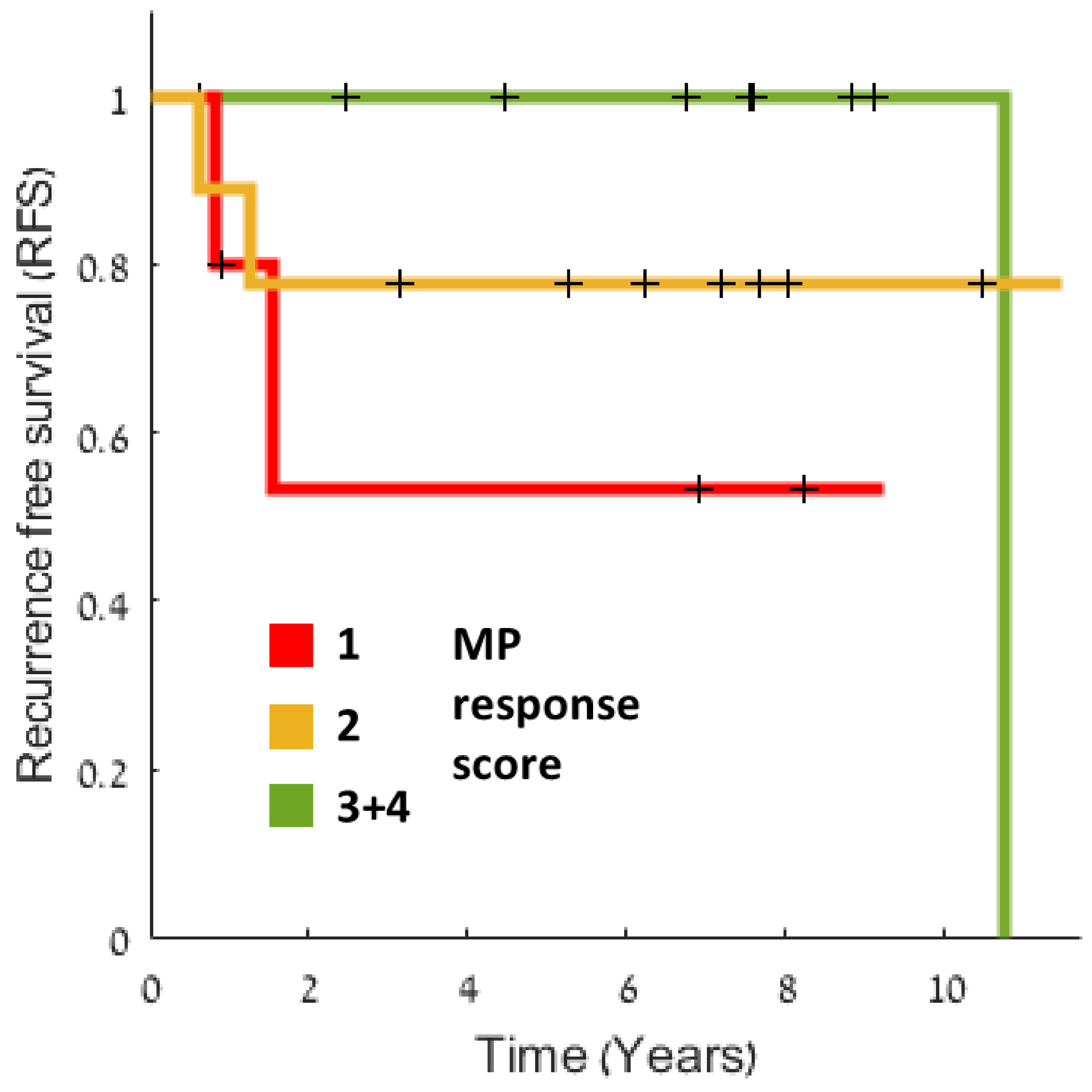


**Figure S7: Kaplan-Meier curve of recurrence free survival for the entire cohort by their MP response score (Log rank p-value 0.16).**

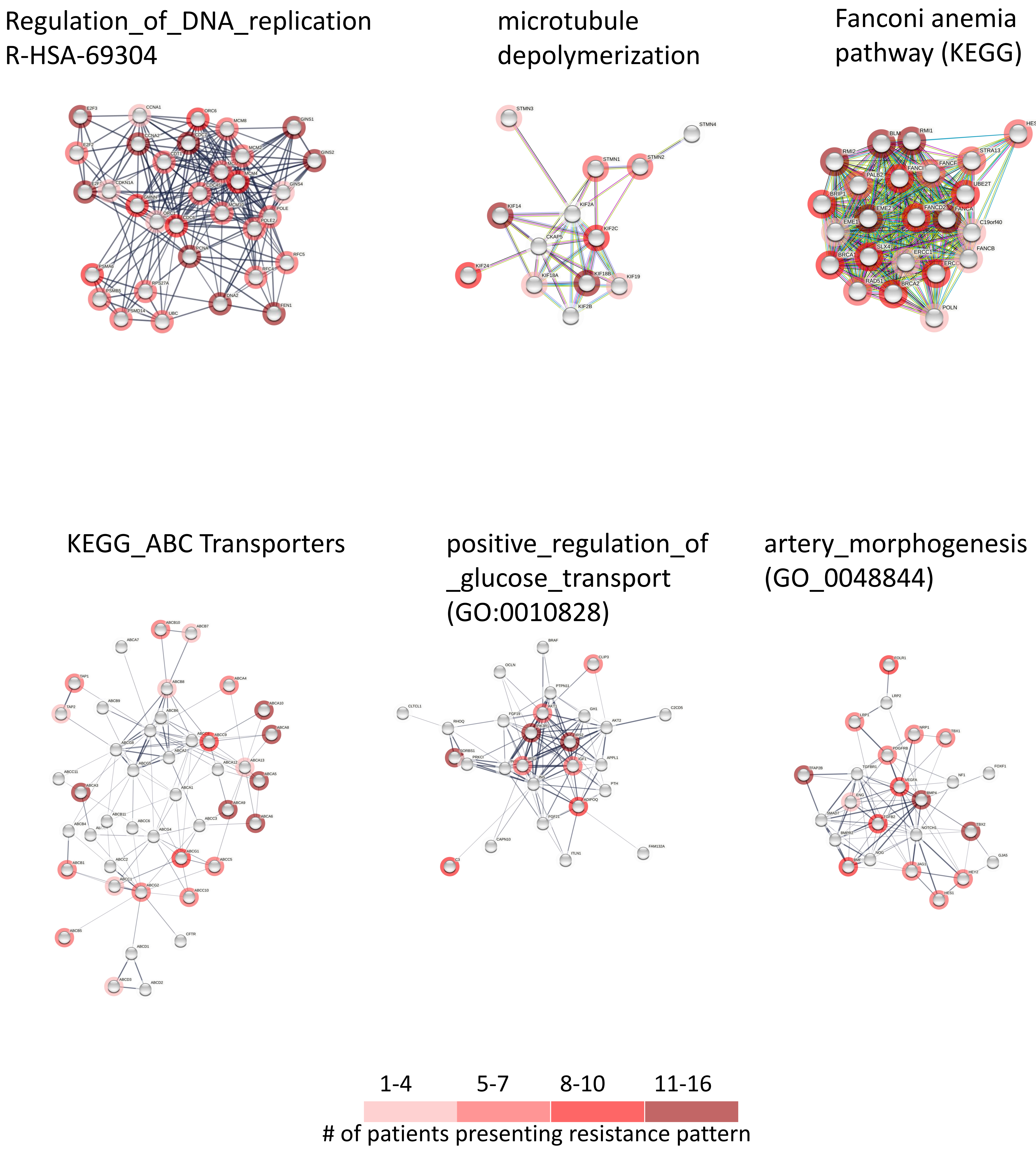


**Figure S8: Hubs of resistant genes in selected dysregulated pathways**. Network representation (STRING) of selected pathways. Genes are colored by the number of genes with resistant patterns.

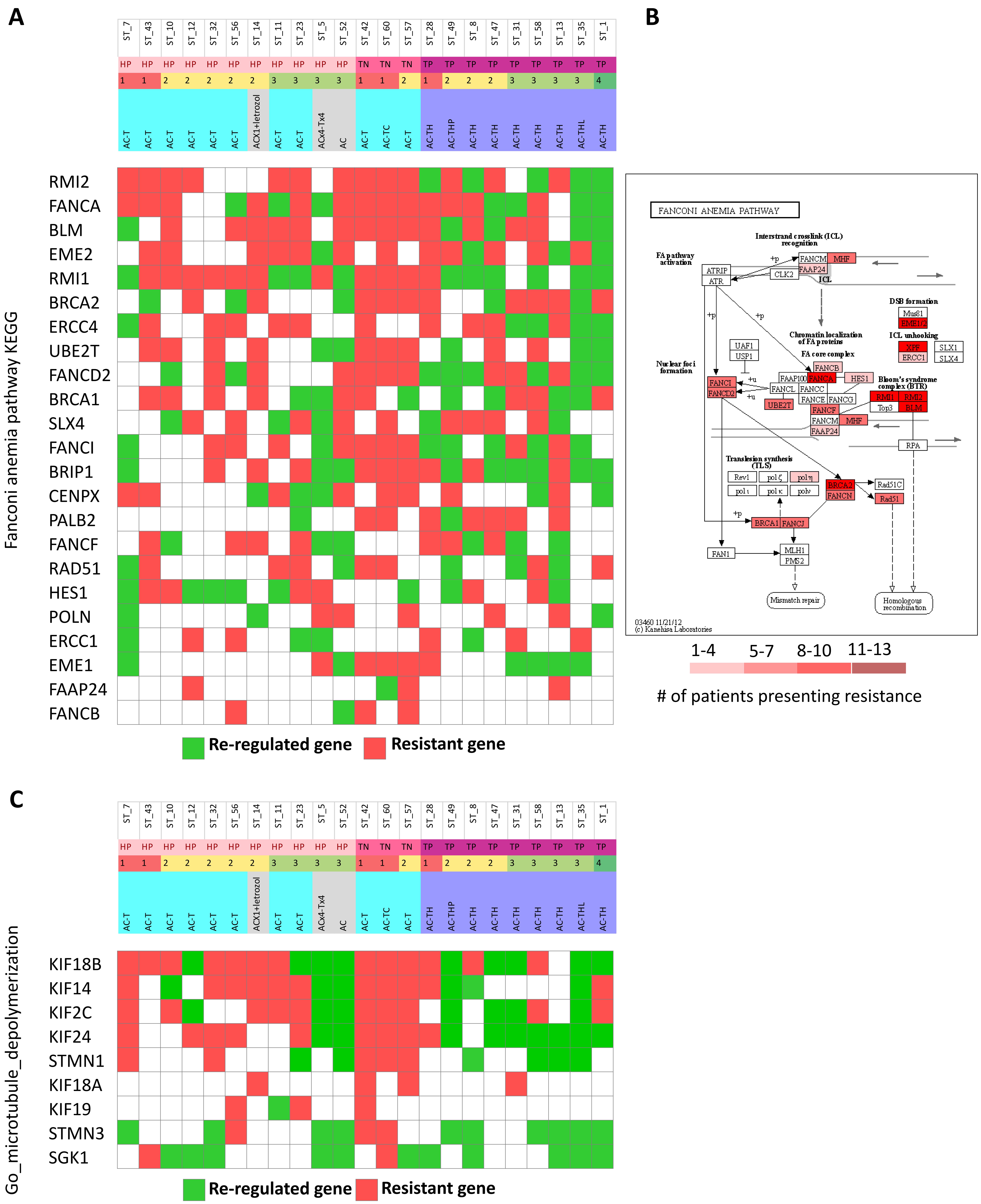


**Figure S9: Heat maps presenting the modes of resistance/reregulation in two representative dysregulated pathways.** A. All differentially expressed genes in the Fanconi Anemia pathway (KEGG) are colored by either the gene pattern was re-regulated (green) or resistant (red). Clinical parameters are presented for each patient, including subtype, response score and administered chemotherapy. B. A scheme of the Fanconi Anemia pathway (KEGG) colored by the number of patients presenting resistance pattern in each gene C. All differentially expressed genes in the Go_microtubule_depolymerization pathway and their mode of resistance or re-regulation in each patient.

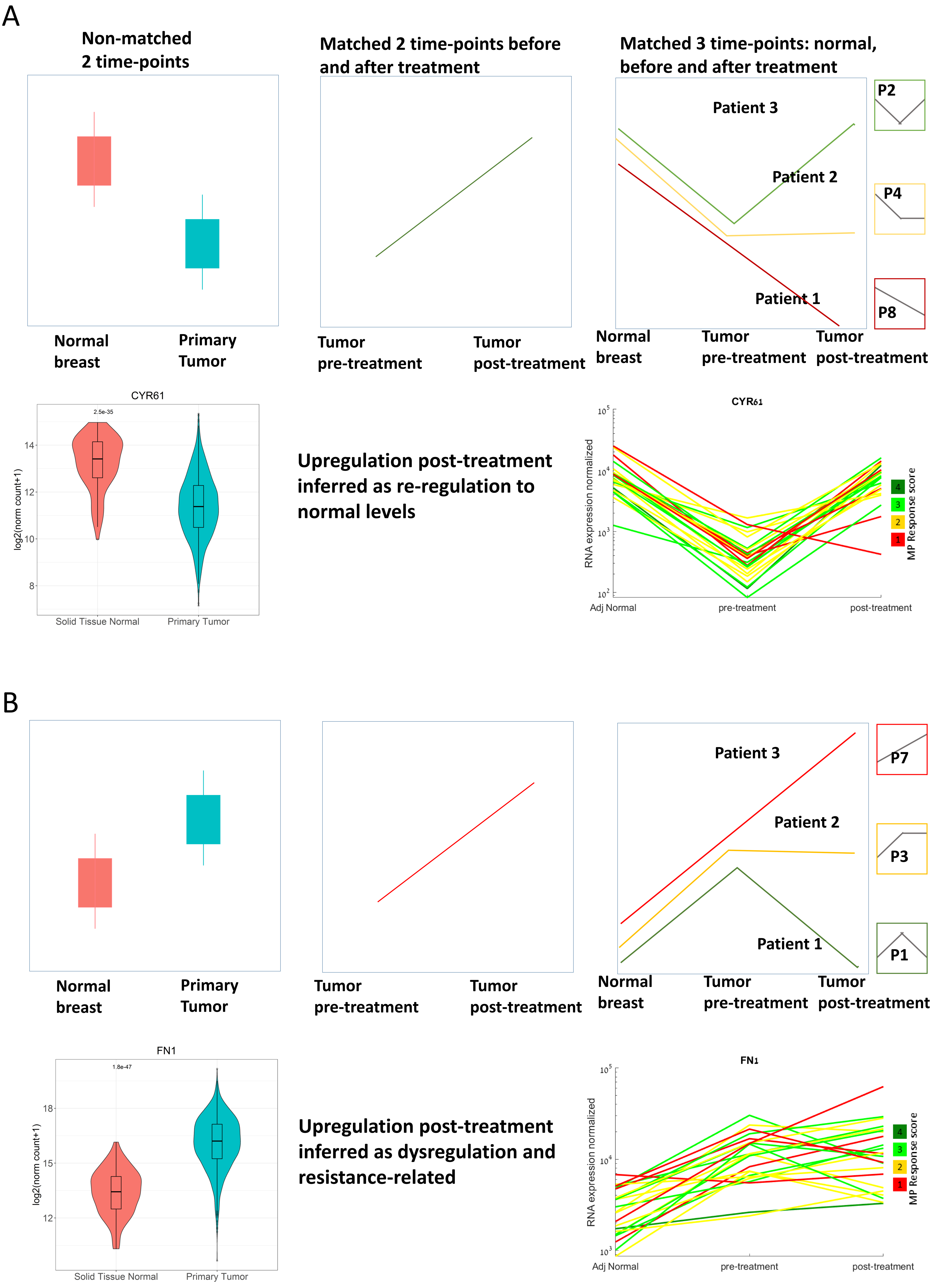


**Figure S10: Resolving resistance by pattern analysis of matched three time points.** Two theoretical scenarios and data examples of genes that are upregulated post treatment, but their normal levels relative to the tumor are opposite. pattern analysis enables to differentiate between interpretation of resistance vs. re-regulation.
